# Lung architecture amplifies tissue deposition in an agent-based model of fibrotic development

**DOI:** 10.1101/2025.04.01.646600

**Authors:** Dylan T. Casey, Alexander Sosa, Vitor Mori, Joseph K. Hall, Béla Suki, Bradford J. Smith, Jason H.T. Bates

## Abstract

Idiopathic pulmonary fibrosis (IPF) is a progressive interstitial lung disease where excessive extracellular matrix (ECM) deposition and remodeling stiffens the lung, impeding its function. Many factors are known to contribute to the development of this fibrosis, but a lack of conclusive understanding endures because of their complex nature. The modification of ECM and the unique architecture of the lung are such factors in IPF’s propagation and not solely casualties. Their effects on fibrogenesis are not known and tricky to study. We apply a computational methodology known as an agent-based model (ABM) to simulate cellular behavior as automata. Our ABM is a tissue maintenance model where agents modify tissue density to sustain a global mean and variance to represent the cyclic turnover of ECM. Agents traverse and interact with high fidelity architecture obtained through micro computed tomography (microCT) of mouse lung tissue. The properties of the ABM are validated to microCT of fibrotic mouse lung tissue. We find that increasing cell density is sufficient for fibrogenesis, but that the lung architecture led to more tissue deposition. Our model suggests that lung structure is a relevant contributor to the pathogenesis of IPF.

---

Idiopathic pulmonary fibrosis (IPF) is a progressive lung disease characterized by a stiffening of tissue through excessive or aberrant tissue deposition^1^. Mechanisms of fibrotic development are complex, varied, and speculative^2^. However, the extracellular matrix (ECM) and its dysfunction remain an essential component of proposed methods of pathogenesis. It is well established that the lung microarchitecture is significantly remodeled in fibrosis^3–6^. Once thought of as a consequence of the condition, the extracellular matrix (ECM) and lung architecture have been shown to play an active role in fibrotic progression^7^. For instance, it has been suggest that the fibrotic ECM creates a positive feedback loop^8^ and that profibrotic cell phenotypes differentiate more frequently on stiffer matrix^9^. Appropriately, using 3D matrix or decellularized lung tissue for in vitro models has become more common, reinforcing the importance of the ECM and architecture^10,11^. However, these mechanisms occur after fibrosis has started to develop or is established. Whether the ECM and lung architecture contribute to fibrogenesis has yet to be determined.

IPF is only detected when the disease has already progressed so there is no certainty as to its early structure. Hence, investigating with a computational model can test likely scenarios and find dynamics essential to the development of fibrosis. Agent-based models (ABM) are particularly useful in biology because they can simulate cellular behaviors via automata called agents. Each agent has a set of rules it obeys such as how to move or when to deposit tissue to mimic complex cellular behaviors. We have previously presented such a model that emulates the cyclic nature of ECM maintenance^12^. However, these models are frequently two-dimensional in the interests of simplicity and ignore the intricacies of their environment’s topology.

In the present study, we developed an ABM that includes a 3D lung architecture for agents to traverse, maintain, and/or modify in the process of creating aberrant ECM. The lung architecture was obtained via ex vivo micro computed tomography (microCT) to capture high resolution parenchymal areas of mouse lung tissue. We validated the model by running agents on tissue matched to control mouse microCT scans and comparing them results from bleomycin-induced fibrosis mouse microCT scans. We also compared the results obtained to those obtained using a cubic lattice to represent a simple tissue structure.

## Results

The cumulative distribution function (CDF) of contrast-enhanced Hounsfield units (ceHU) from bleomycin-induced mice were skewed to higher values relative to control mice in Fig. 1. They had means 2-3 times higher and the variation in bleomycin-induced mice was large with one sample overlapping with the control mice. Control mice ceHU were quite similar to each other. The CDF of the lung parenchymal subsections (Fig. 1a) of all mice were nearly identical to the CDF of the whole lung scan they came from (Fig. 1b).

**Figure 1:**
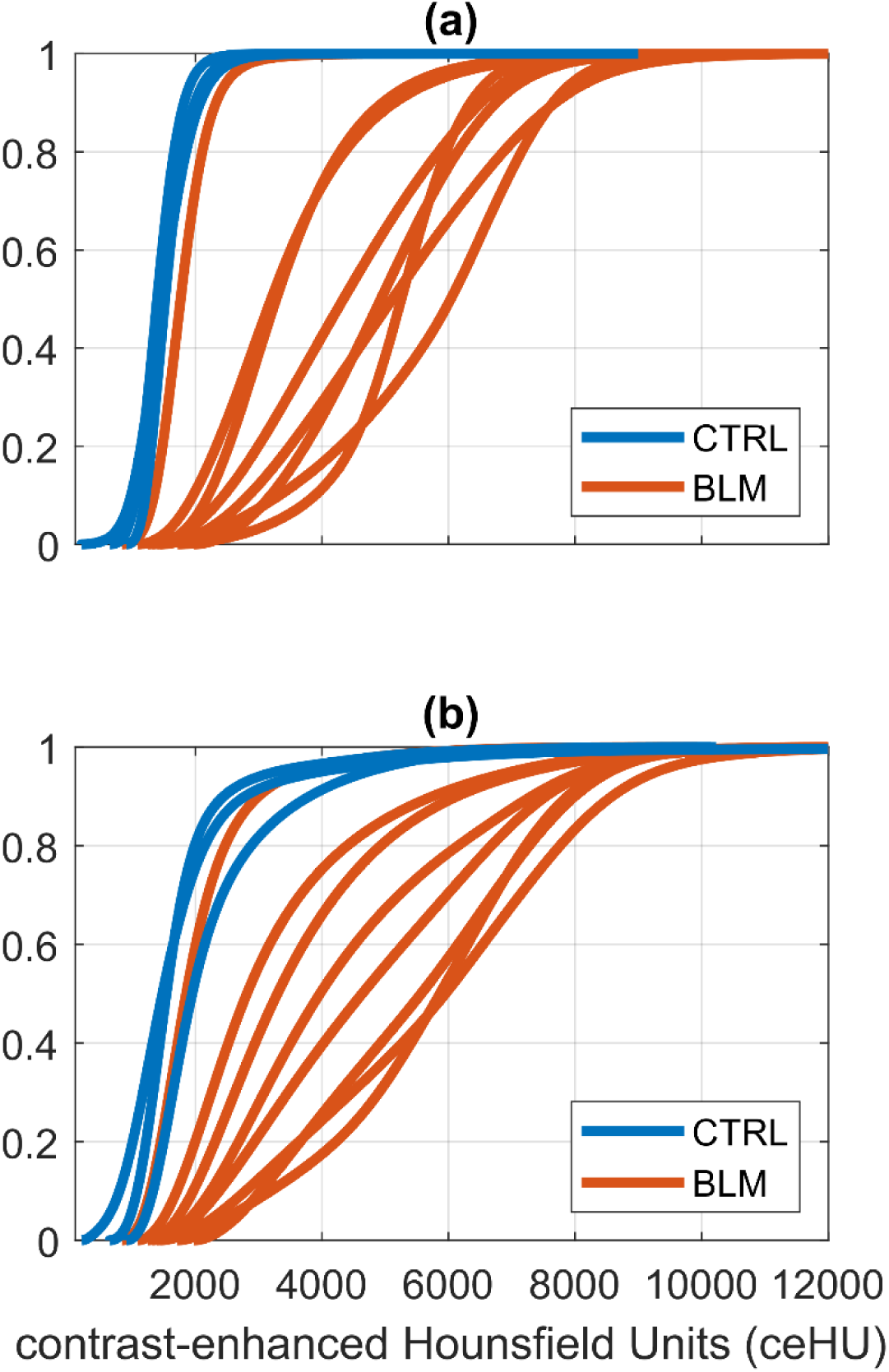
Cumulative distribution functions (CDF) for control and bleomycin-induced fibrosis mice of (a) parenchymal subsections and (b) whole lung slices.

The CCDF of volumetric slices of airspaces in lung parenchymal subsection in Fig. 2a were best fit with the sum of two exponential functions whose coefficients are in Table 2. The volume tissue fraction for the parenchymal cuboids were not different (P=0.7040). In general, the bleomycin-induced mice had larger airspaces. For the whole lung slices, the CCDF (Fig. 2b) took on a sigmoidal shape due to large airspace slices being more prominent and having more volume. The bleomycin-induced mice maintained the trend of having larger airspaces than the control mice. We observed a decreased lung volume capacity at the maximum physiologically relevant pressure of 30H_2_O, as well as an increase in variation of lung volumes (Fig. 3). Lung mechanics measurements demonstrated stiffer lungs and greater tissue dampening at all PEEP values (Supplement Fig. 2). Hysteresivity and airway impedance were relatively unchanged, but impedance was slightly elevated for bleomycin-induced mice at low PEEP (Supplement Fig. 3).

**Figure 2:**
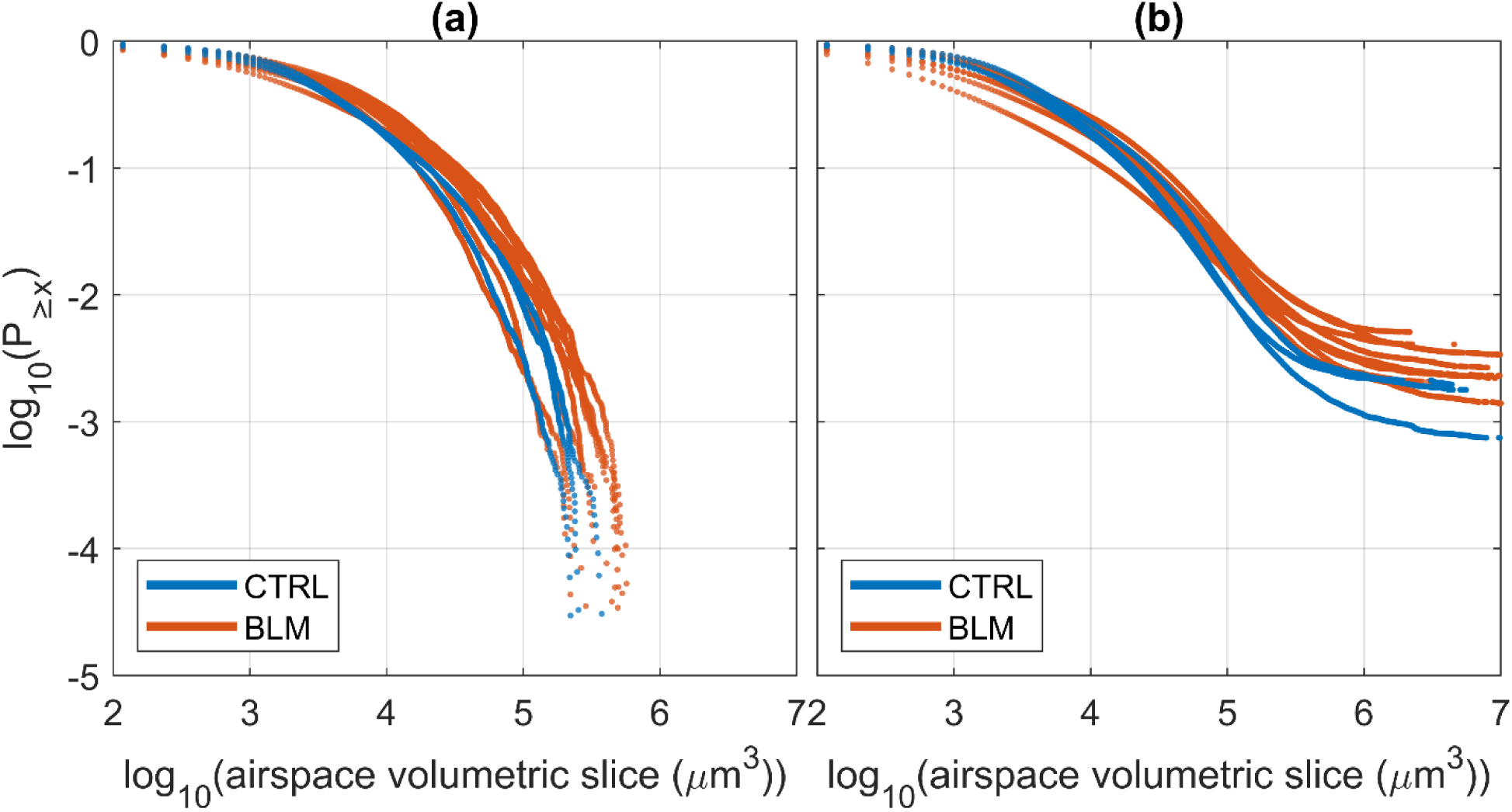
Complementary cumulative distribution functions (CCDF) of cross-sectional airspace sizes aggregated from the x-, y-, and z-directions for **(a)** parenchymal subsections and **(b)** whole lung slices. Coefficients for an exponential fit in (a) can be found in Table 2.

**Figure 3:**
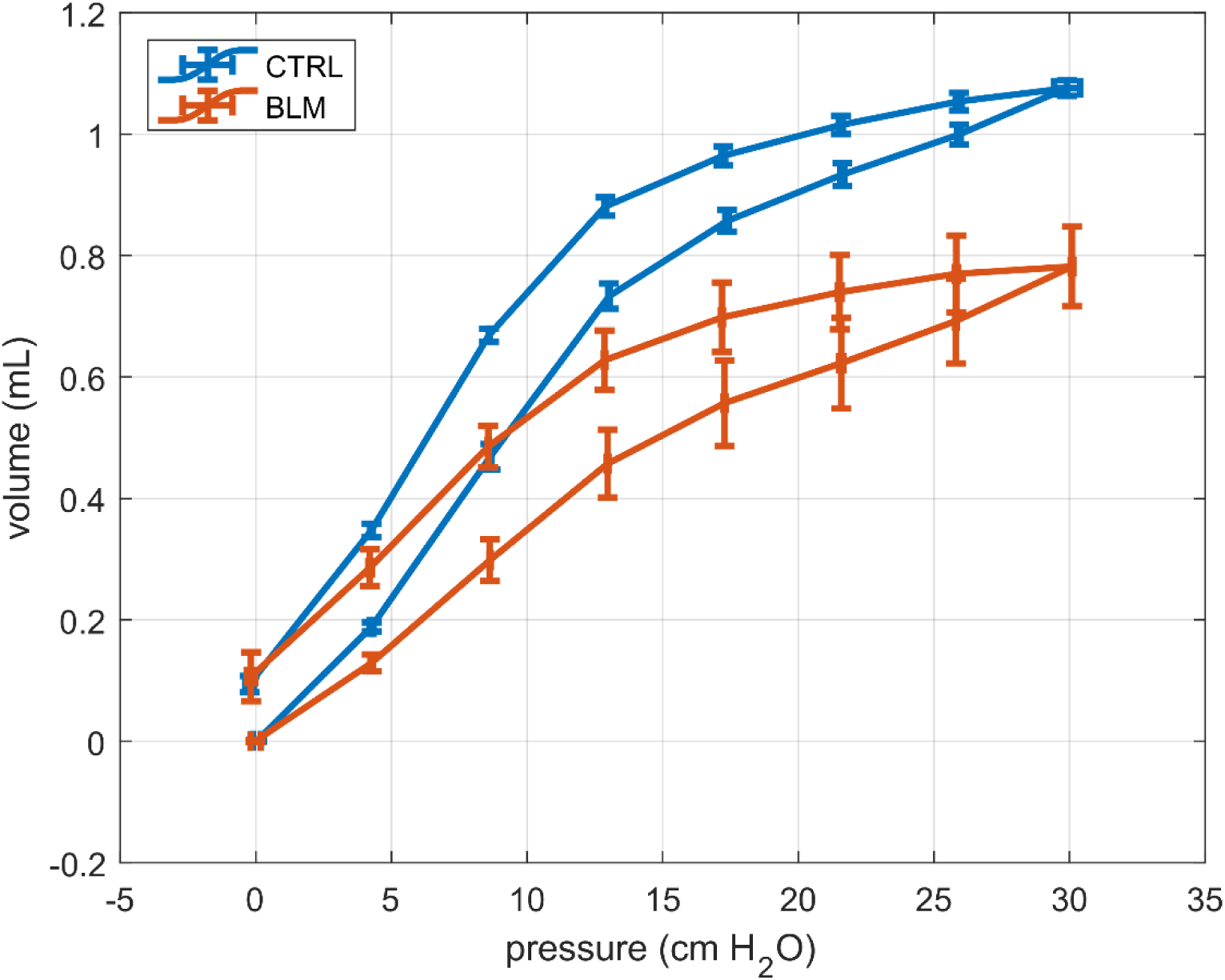
Pressure-volume curves of control (n=3) and bleomycin-induced fibrosis mice (n=4)

**Table 1:** Mouse records of lung volume and weight.

|  | Lung volume (mL) | Starting weight (g) | Harvest weight (g) |
| --- | --- | --- | --- |
| CTRL 1 | 0.91 | 30.3 | 30.3 |
| CTRL 2 | 0.84 | 30.1 | 30.1 |
| CTRL 3 | 0.79 | 31 | 31 |
| BLM 1 | 0.68 | 26.6 | 27 |
| BLM 2 | 0.73 | 25.8 | 23.4 |
| BLM 3 | 0.7 | 24.5 | 23.9 |
| BLM 4 | 0.66 | 26.1 | 23 |

**Table 2:** Volume fraction of parenchymal subsection samples. Exponential fitting parameters to CCDF by lung sample in Figure 2a.

|  | Volume fraction | a | b | c | d | R2 |
| --- | --- | --- | --- | --- | --- | --- |
| CTRL 1 | 0.6826 | 0.7078 | -0.0003628 | 0.2254 | -0.00003875 | 0.9985 |
| CTRL 2 | 0.7884 | 0.5929 | -0.0003748 | 0.2851 | -0.00006013 | 0.9988 |
| CTRL 3 | 0.7119 | 0.7415 | -0.0003393 | 0.2262 | -0.00003891 | 0.9992 |
| BLM 1.1 | 0.4725 | 0.4876 | -0.0002926 | 0.2987 | -0.00002943 | 0.9947 |
| BLM 1.2 | 0.7018 | 0.6202 | -0.0003128 | 0.3047 | -0.00004016 | 0.9985 |
| BLM 2.1 | 0.7428 | 0.509 | -0.0002484 | 0.3727 | -0.00004188 | 0.9974 |
| BLM 2.2 | 0.7438 | 0.522 | -0.0003312 | 0.3867 | -0.00004783 | 0.9968 |
| BLM 3.1 | 0.6918 | 0.4579 | -0.0002981 | 0.3011 | -0.00003273 | 0.9936 |
| BLM 3.2 | 0.7895 | 0.5101 | -0.000453 | 0.4047 | -0.00008079 | 0.9988 |
| BLM 4.1 | 0.7903 | 0.5589 | -0.0003443 | 0.3634 | -0.00005959 | 0.9986 |
| BLM 4.2 | 0.6847 | 0.5271 | -0.0004155 | 0.2475 | -0.00003401 | 0.9927 |

Figure 4 displays the means and variances of the model that ran on the parenchymal subsections partitioned into 128 27 x 27 x 27 voxel cubes over 10,000 timesteps. The mean of the ABM matched was marginally greater than that of the model at baseline (*β*), when cell density was 13.65% with both lung and lattice architectures (Fig. 4a). Both variances were low and constant, although greater in the lattice architecture (Fig. 4d). At a cell density of 17.5%, the means increased to near twice the baseline with the lattice architecture having a lower mean than the lung architecture (Fig. 4b). There was very little overlap in their 90% confidence intervals. The variances were elevated by 4-fold, although the lattice architecture had a greater mean variance and a greater spread (Fig. 4e). At 20% cell density, The difference between the means of the lung and lattice architecture increased, while the means themselves increased to ∼3β (Fig. 4c). Variances increased an average of 10 fold above baseline, with that of the lattice architecture being greater than the lung architecture (Fig. 4f).

**Figure 4:**
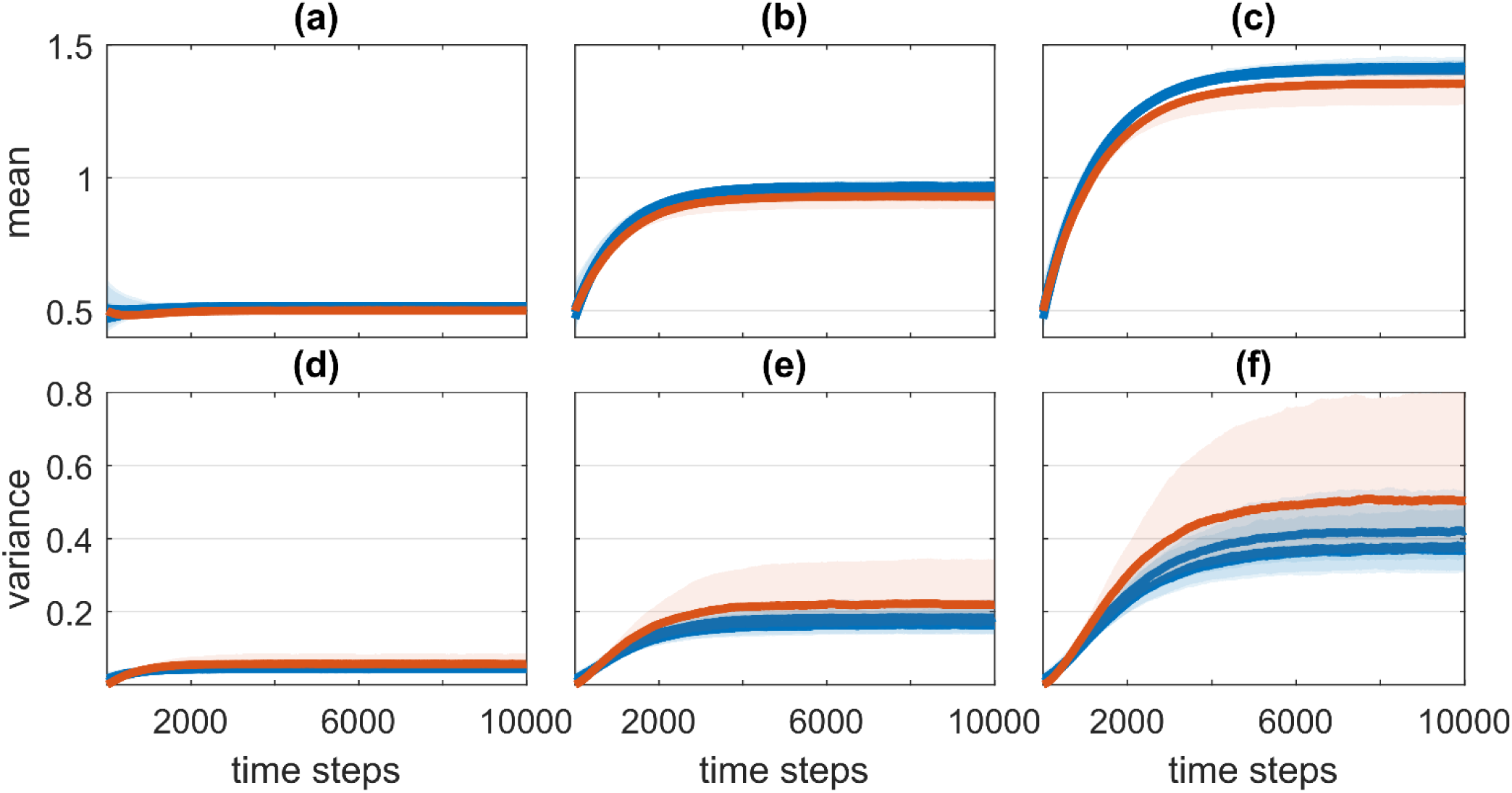
Mean and variances of ABM tissue remodeling over 10,000-time steps for lung and lattice architectures. Left column (a, d) represents theresults fromagent density of 13.65%, center column (b, e) are from agent density of 17.5%, and right column (c, f) are from agent density of 20%. Solid line represents the median and the shaded region surrounding it represents the 90% confidence interval of the 128 partitioned cubes.

Figure 5 displays examples of cuboid parenchymal sections from a control mouse used as architecture for the ABM. The original structure with its scanned ceHU is shown in Figure 5a for comparison. The ABM results are shown at time step 10,000 for cell densities of 13.65% (Fig 5b), 17.5% (Fig 5c), and 20% (Fig 5d). The ceHUs in Fig. 5b are roughly the same as in Fig. 5a and are 2 and 3 times greater in Figs. 5c and d, respectively.

**Figure 5:**
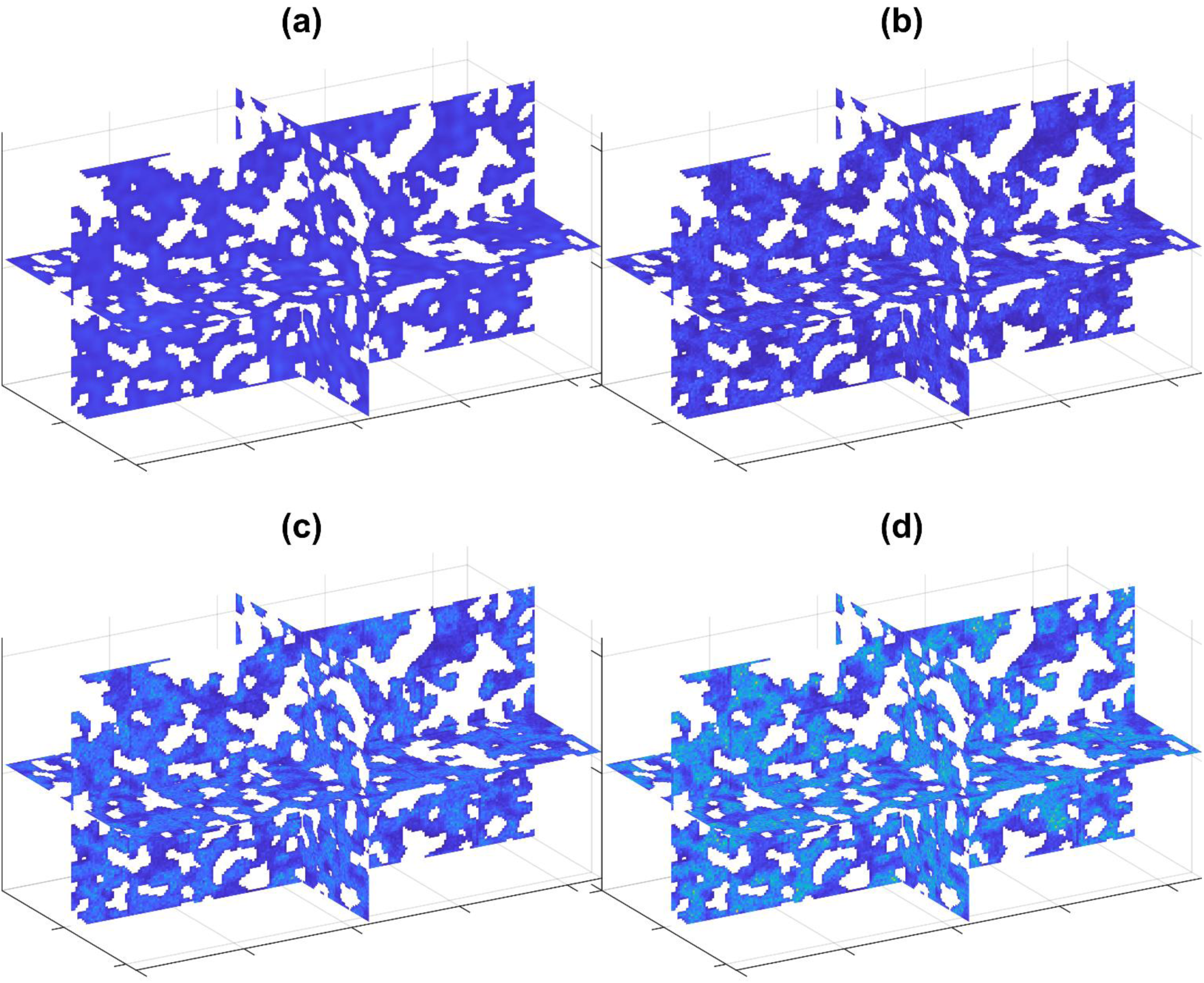
Cross-sectional slices planes of a parenchymal subsection from a control mouse. Each structure is a 108 x 108 x 216 voxel cuboid that was partitioned into 128 27 x 27 x 27 voxel cubes for the purpose of running the ABM. Original structure from microCT is in (a). Results of tissue remodeling from the model running for agent density (b) 13.65%, (c) 17.5%, and (d) 20%. Color represents the ceHUwith all plots on the same scale, with white corresponding to empty space i.e. airspace.

After converting the ABM to the ceHU scale, we found that, at a cell density of 13.65%, the model maintained its distribution of tissue density on both architectures (Fig. 6a). The CDF at cell densities of 17.5% and 20% overlapped with the CDF from the scanned BLM mice lungs (Figs. 6b, c). However, the lattice architecture distribution produced more low-density tissue as cell density increased.

**Figure 6:**
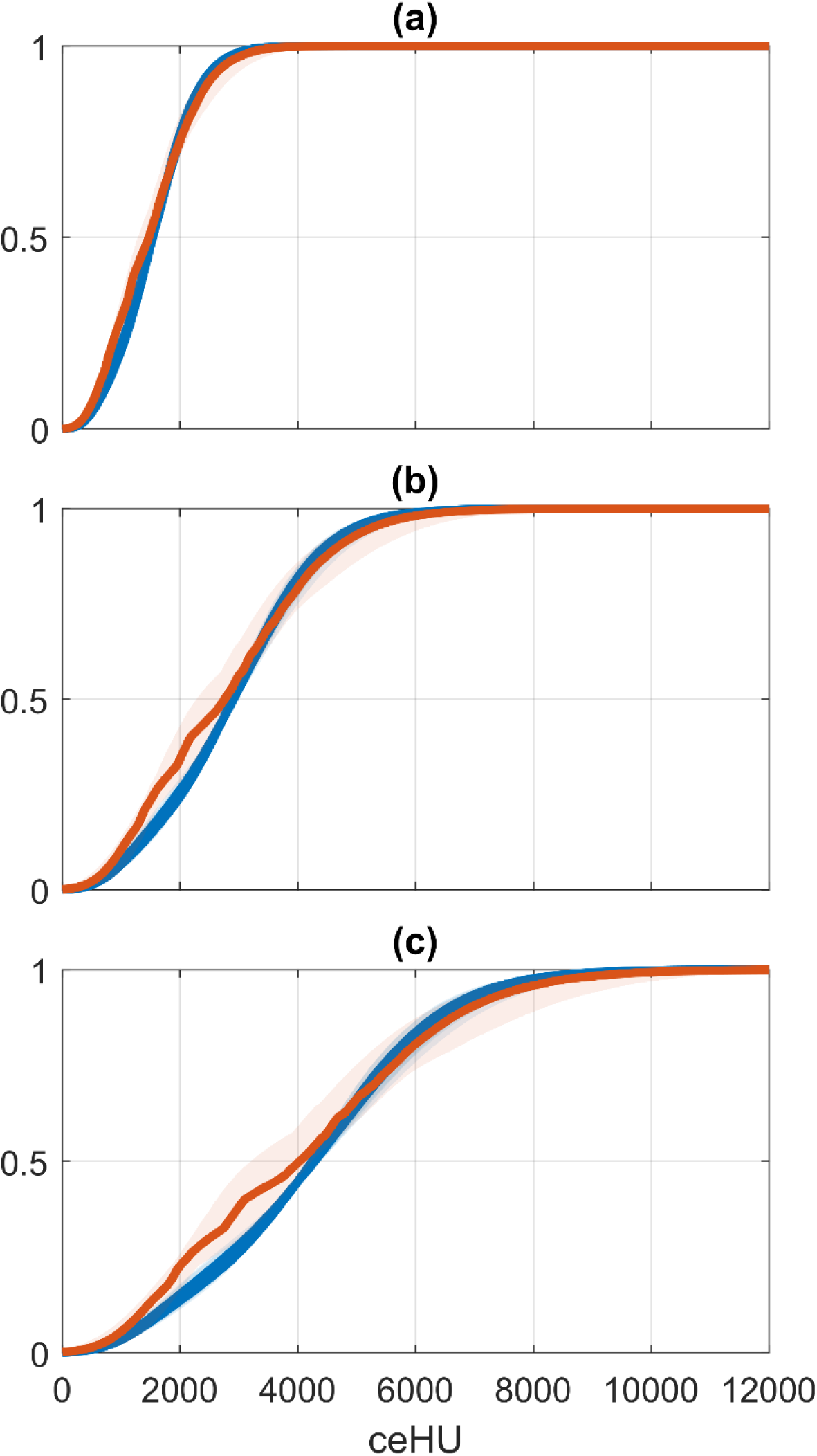
CDF of results from ABM tissue remodeling for lung and lattice architectures. Results are from an agent density of (a) 13.65%, (b) 17.5%, and (c) 20%. Solid line represents the median and the shaded region surrounding it represents the 90% confidence interval of the 128 partitioned cubes.

As a function of partitioned cube volume, the means at time step 10,000 decreased slightly in a linear fashion with increasing volume (Fig. 7a). This effect was exacerbated with a lower slope (Table 3) in the lung architecture and as cell density increased. The variance also decreased linearly with volume (Fig 7b), however, the slope decreased with cell density and not architecture (Table 4). The fraction of unique locations that agents visited over 10,000 time steps decreased with increasing volume, exhibiting a power law relationship (Fig. 7c). Agents on the lattice architecture visited fewer unique voxels for all volumes, although the exponent of the power law relationship was little different from that obtained when the agents were on the lung architecture (Table 5).

**Figure 7:**
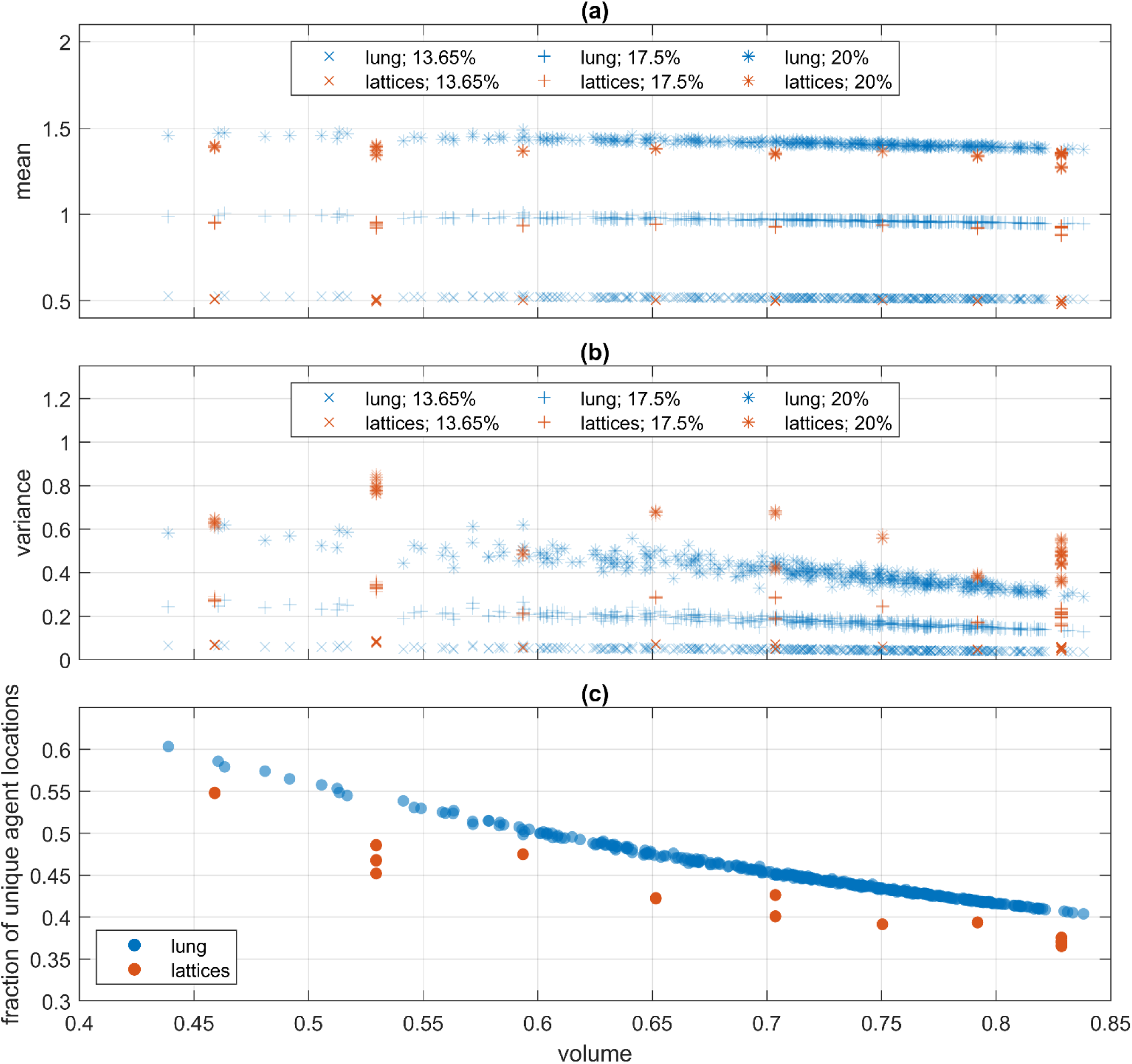
Model metrics at time step 10,000 as a function of volume. **(a)** Volume vs mean vs of each partition for each agent density in lung and lattice architecture. **(b)** Volume vs variance of each partition for each agent density in lung and lattice architecture. Linear regression coefficients for data in (a) and (b) are in Table 3 and Table 4, respectively. **(c)** Volume vs mean fraction of unique voxel locations agent visits for lung and lattice architectures. Data from all agent densities are represented in (c) and power-law fits for data are in Table 5.

**Table 3:**
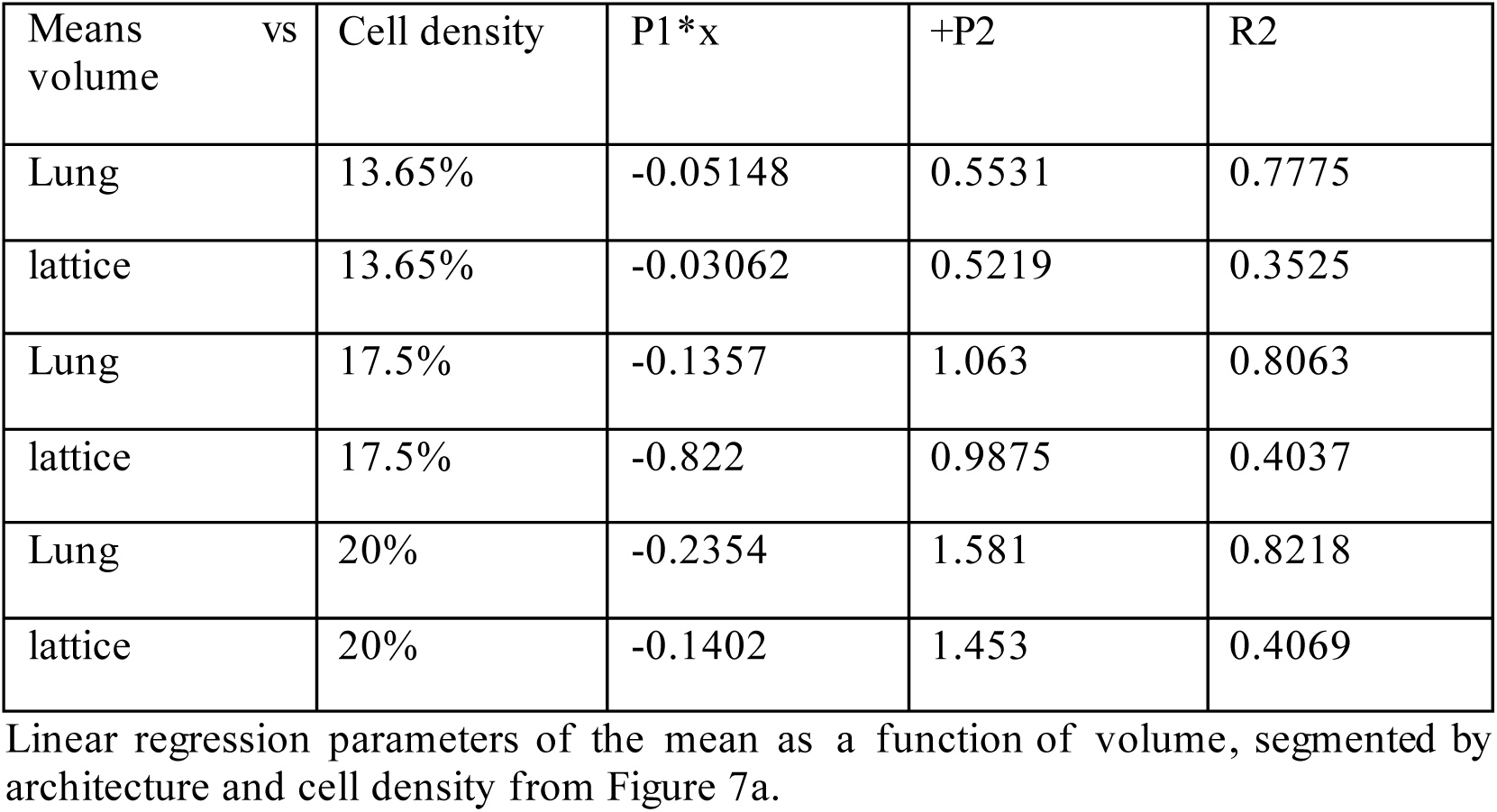
Linear regression parameters of the mean as a function of volume, segmented by architecture and cell density from Figure 7a.

**Table 4:** Linear regression parameters of the variance as a function of volume, segmented by architecture and cell density from Figure 7b.

| Variance vs vol | Cell density | P1*x | +P2 | R2 |
| --- | --- | --- | --- | --- |
| Lung | 13.65% | -0.0709 | 0.09679 | 0.8071 |
| lattice | 13.65% | -0.07412 | 0.1131 | 0.6119 |
| Lung | 17.5% | -0.3168 | 0.4027 | 0.8171 |
| lattice | 17.5% | -0.3336 | 0.4741 | 0.6014 |
| Lung | 20% | -0.7732 | 0.9546 | 0.8210 |
| lattice | 20% | -0.8138 | 1.127 | 0.5826 |

**Table 5:** Power-law fit parameters of the mean probability of a voxel visit as a function of volume, segmented by architecture from Figure 7c.

|  | a | x^b | R2 |
| --- | --- | --- | --- |
| lung | 0.362 | -0.6329 | 0.9977 |
| Lattice | 0.3311 | -0.609 | 0.9509 |

Voxel connectedness, defined as the number of adjacent voxel it touches either by face, edge, or corner, were distributed differently between the lung and lattice architectures. Both architectures had large amounts of fully connected voxels, meaning they were on the interior of the matrix away from the airspaces (Figs. 8a, b). The natural architecture had voxels of every connectedness, distributed in a decreasing exponential fashion with connectivity (Fig. 8a). The lowest connectivity value for the lattice architecture was 8, but most were 18 or greater (Fig. 8b). The probability that a voxel was visited in the lung architecture model increased linearly with connectedness, with variation also increasing with connectedness (Fig. 8c). The pattern for lattice architecture visitation did not exhibit a clear pattern but variances were large (Fig. 8d). All of these behaviors were independent of agent density.

**Figure 8:**
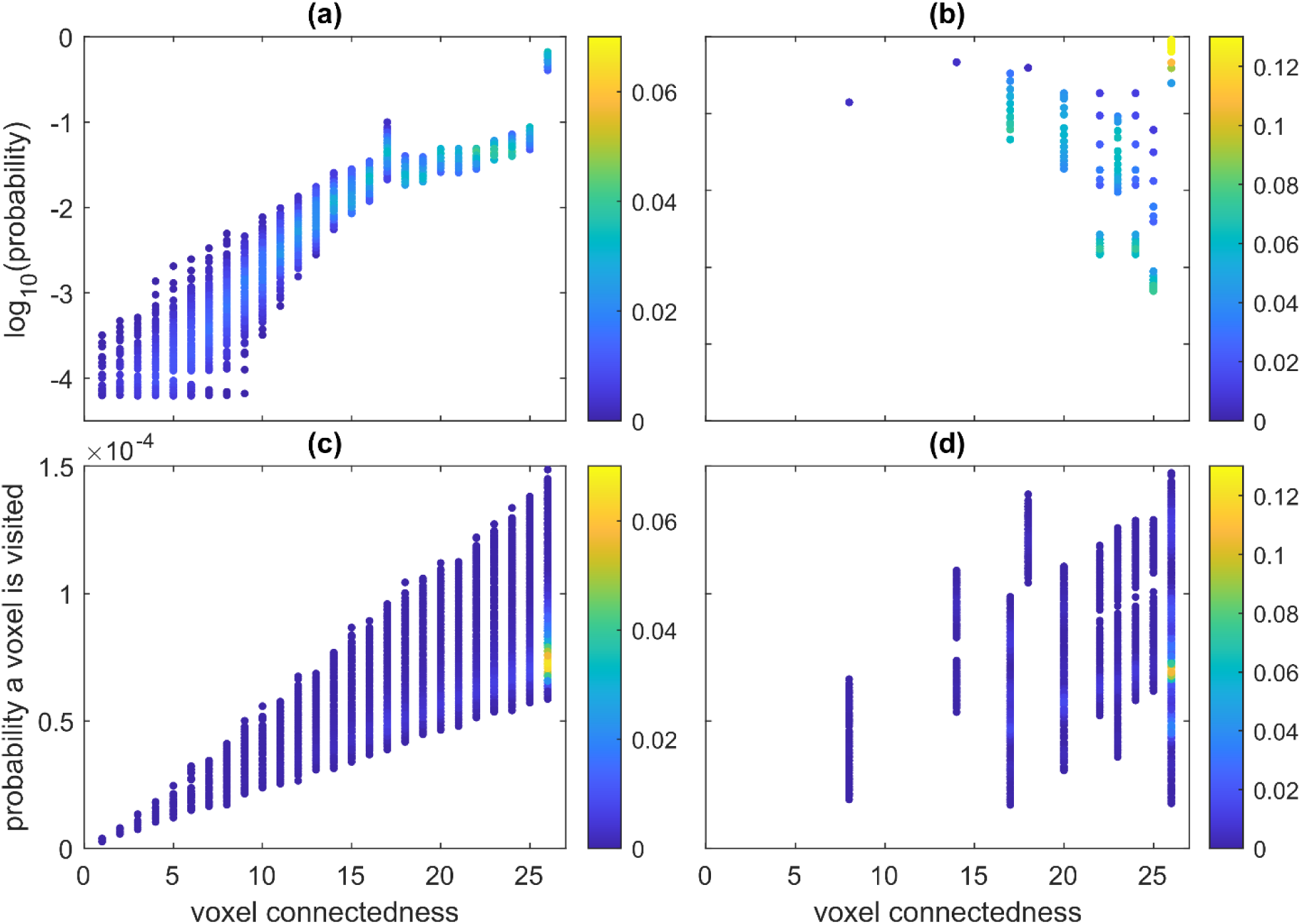
Probability mass function of voxel connectedness in (a) lung architecture and (b) lattice architectures. Probability a voxel is visited as a function of its connectedness for (c) lung architecture and (d) lattice architectures. Color represents density.

Voxel connectedness influenced the tissue density in the ABM shown in Fig. 9. Higher connectedness correlated with higher ceHU in the lung architecture model at baseline cell density (Fig. 9a) and expanded nonlinearly with 17.5% and 20% cell densities (Figs. 9b, c). Voxels with high connectivity did not have low tissue values and had a minimum near the starting mean in the 20% cell density simulations (Fig. 9c). Fully connected voxels had an increased spread in tissue values as cell density increased. In the lattice architecture model, tissue density increased with the number of agents, but the connectedness had no obvious effect (Figs. 9d-f). Notably, fully connected voxels had the lowest ceHU, and voxels with a connectedness of 18 had a spread in tissue density values that rivaled the fully connected voxels (Fig. 9f).

**Figure 9:**
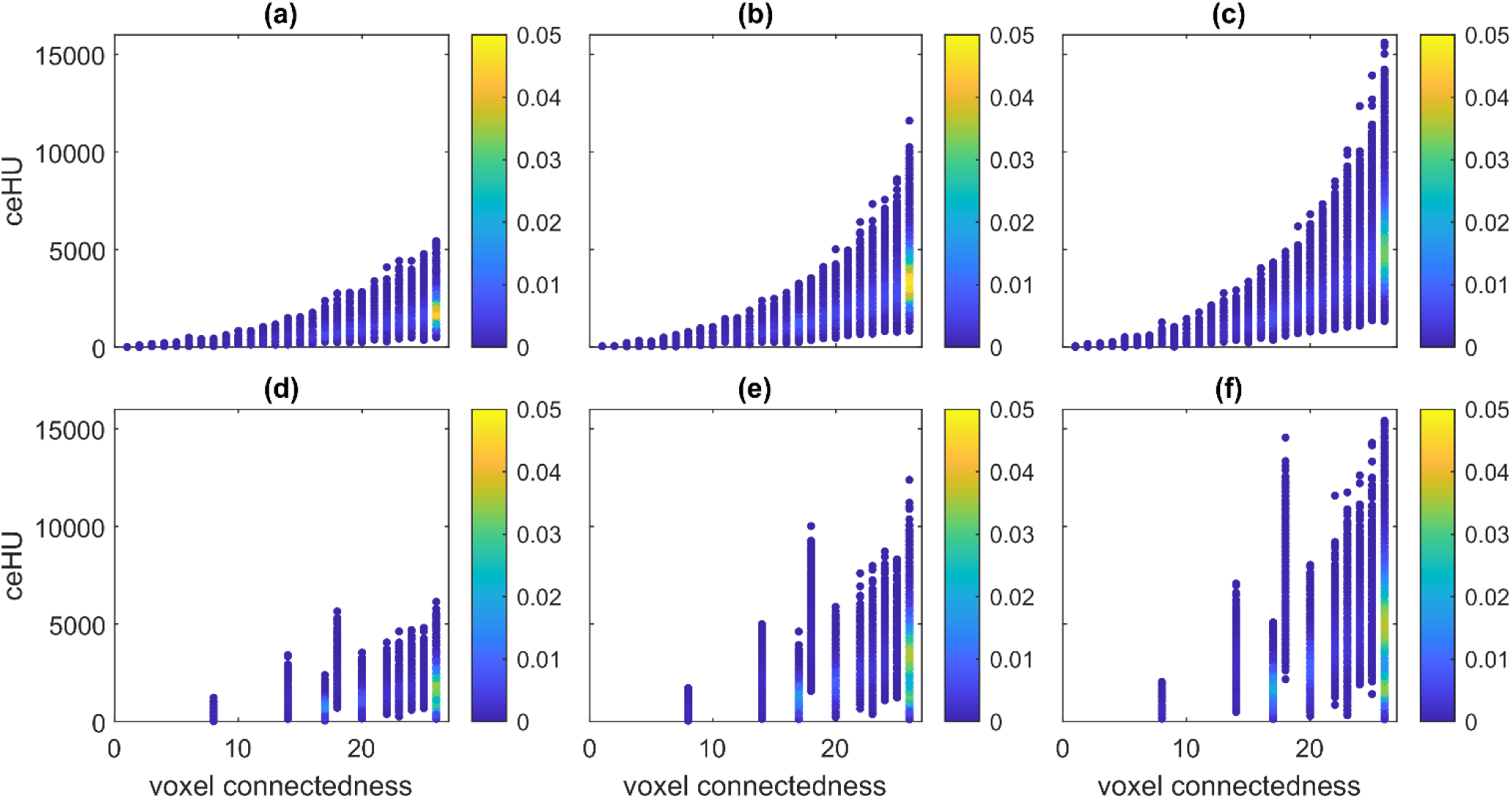
Voxel connectedness vs ceHU of the voxels at time step 10,000 for lung (top row) and lattice (bottom row) architectures. Left column (a, d) represents the results from agent density of 13.65%, center column (b, e) are from agent density of 17.5%, and right column (c, f) are from agent density of 20%. Color represents density.

The most visited voxels were not the ones with the highest tissue densities (Fig. 10). For the lung architecture model, tissue density grew nonlinearly with the probability of being visited (Figs. 10a-c). Highly visited voxels did not have low tissue values, but voxels that had visit numbers around the median were the ones that were concentrated around the mean tissue density. These voxels were also those with the highest tissue densities. The simple lattice architecture model exhibited the same behavior, although at high probability, tissue density increased (Figs. 10 d-f). Three distinct clusters formed based on the probability of being visited. These clusters expanded to higher tissue densities with increasing cell density.

**Figure 10:**
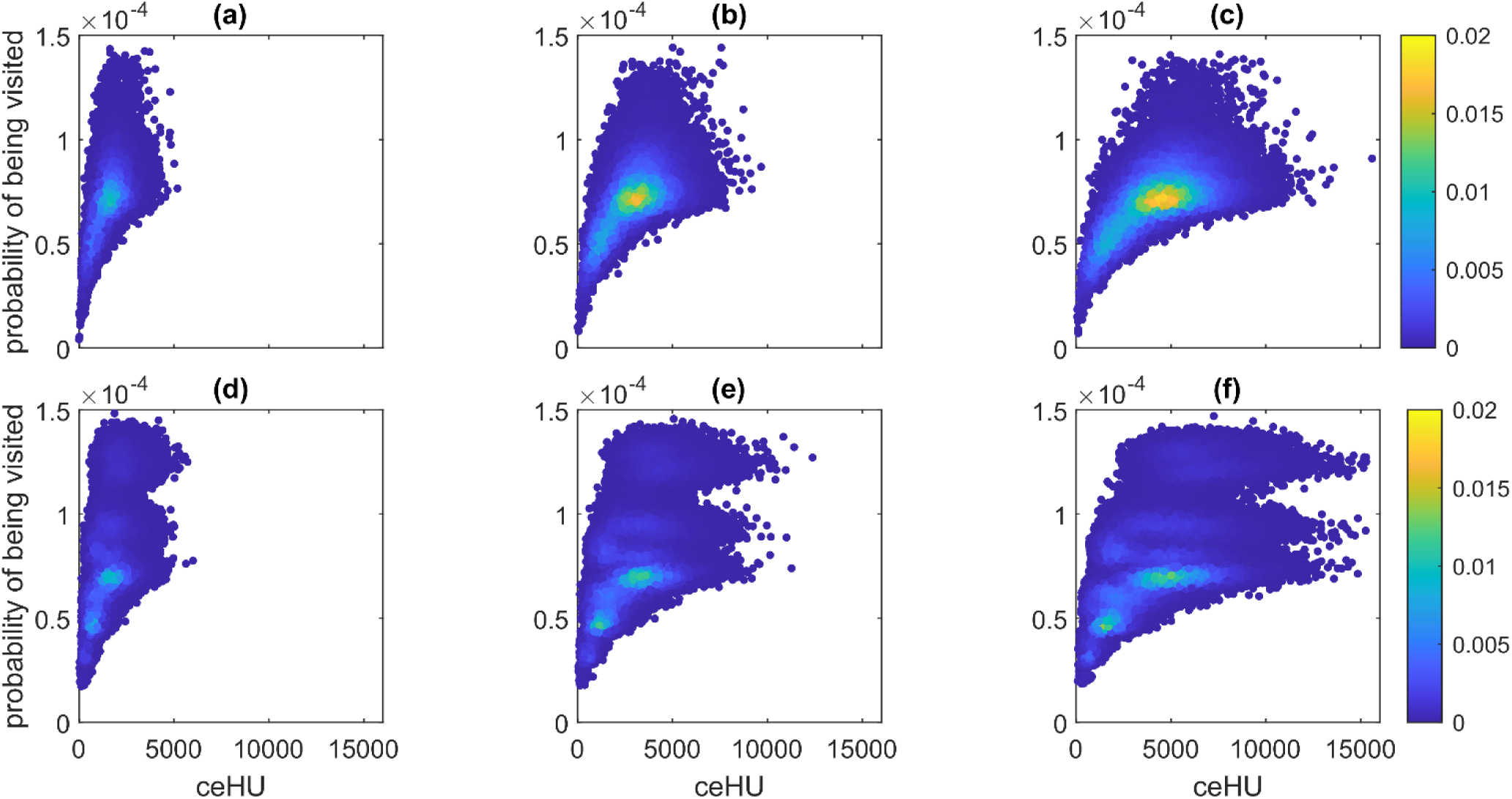
Probability of voxel visits as a function of ceHU at time step 10,000 for lung (top row) and lattice (bottom row) architectures. Left column (a, d) represents the results from agent density of 13.65%, center column (b, e) are from agent density of 17.5%, and right column (c, f) are from agent density of 20%. Color represents density.

Figure 11 shows a positive correlation between models of different cell densities on the same architectures. Increases in tissue density occurred in the same regions, as can be observed in Fig. 5.

**Figure 11:**
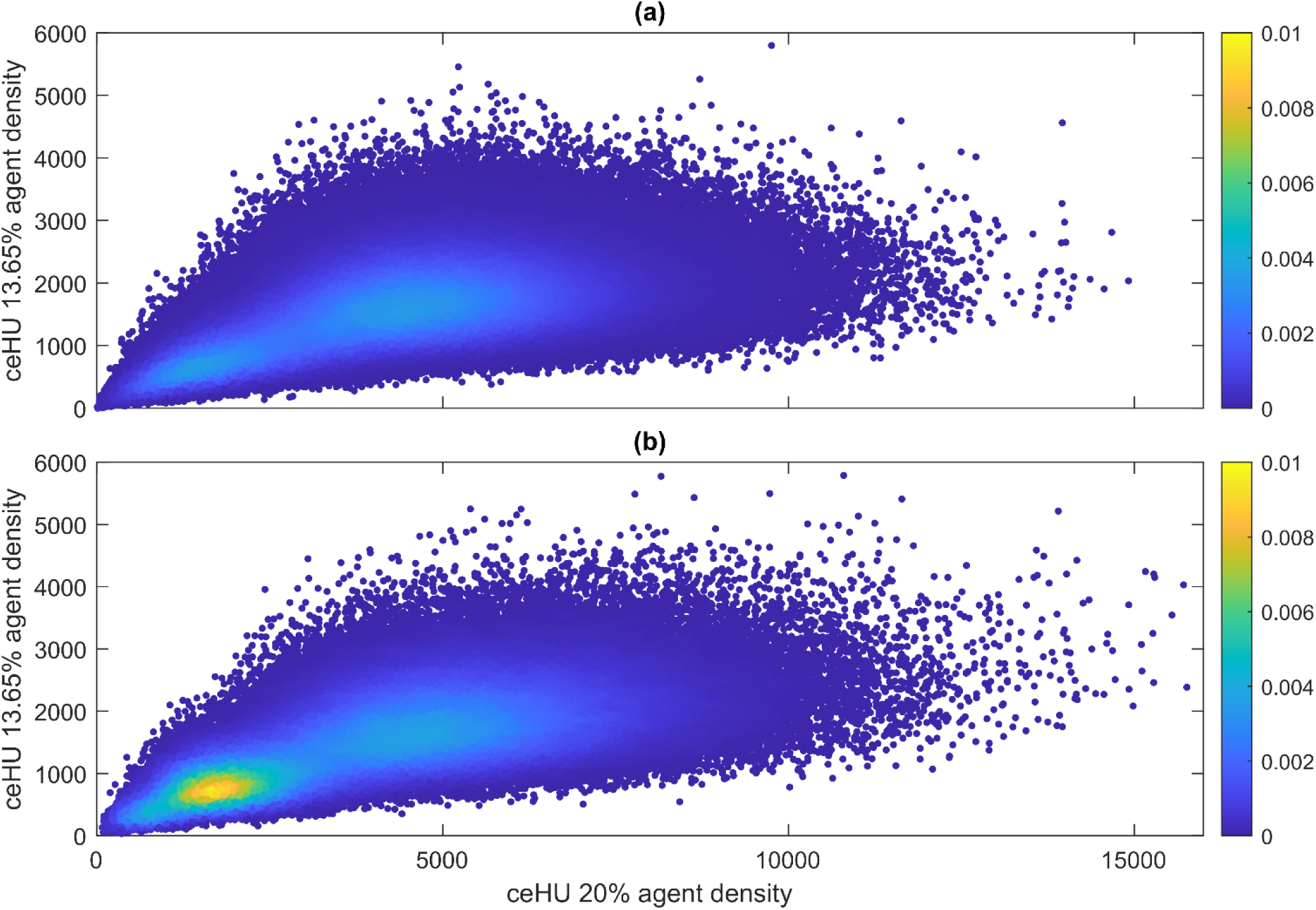
Comparison of ceHUat time step 10,000 between the 13.65% and 20% agent density ABM for (a) lung architecture and (b) lattice architecture. Color bar represents density.

## Discussion

Our agent-based model recapitulates the development of fibrosis showing that a 50% increase in cell density is sufficient to match the tissue density distribution from mice with bleomycin-induced fibrosis. Fibrosis is further exacerbated by the lung architecture itself through its effect on agent movement.

For a given tissue architecture, any random initialization of the model will build up and degrade tissue in essentially the same locations (Fig. 5). Agents move completely randomly in a spatially uncorrelated manner, so the only external influence on agent behavior can be the lung architecture^13,14^. Cell movement, however, is known to be biased even in the absence of external signals^15^. Hence, cell movement is typically modeled as a correlated or persistent random walk meaning previous steps affect the direction of the next step^14^. Cell migration is tightly regulated by a vast set of physiochemical mechanisms, not limited to gradients from chemotaxis and mechanotransduction^16,17^. Cells also move nonrandomly through 3D environments because of the complex organization of the ECM that impedes cell movement or requires remodeling^18,19^. In the absence of information about local tissue biomechanical properties, agent movement is linearly correlated to the connectedness of a given destination (Fig. 8c), despite the connectedness of the voxels being highly skewed (Fig. 8a). Therefore, lower connected voxels are visited disproportionately more often, and the fully connected voxels are visited relatively less frequently. These low-connected voxels may be needed to accurately recreate the tissue density distribution because they consistently have low tissue densities (Figs. 9a-c).

A realistic lung architecture predisposes to the accumulation of ECM compared to an artificial cubic lattice architecture. This may be because, in the former, agents visit more unique locations, which become rarer with deceasing tissue volume (Fig. 7c). A decay in tissue density only occurs when a location is not visited by profibrotic cells within a fixed number of time steps as a result of ongoing degradation by matrix metalloproteinases (MMPs). Thus, the increased motility seen in the lung matrix may inhibit the homeostatic response to a buildup of excessive tissue. In Fig. 10, we can see in both architectures that the voxels with the highest tissue density are not the most visited and thus not do not experience the most deposition of tissue. Also, sudden large deposits of tissue were subject to decay. Persistence is therefore required in our model for fibrotic lesions to develop, which becomes more likely with an increase in cell density. The model predicts that, in order to combat this, regions with higher cell densities are protected from having an increase in the amount of tissue that is laid down while simultaneously being degraded. The drive force behind the cell increase seen in IPF may be fibroblastic foci, which are locations of proliferating fibroblasts unique to IPF, which have a characteristic spatial distribution that may amplify the development of IPF^20^. A very modest increase of cell numbers has been shown in BAL at 21 days in the bleomycin mouse model^21^.

It was necessary to use a contrast agent to exaggerate the normally subtle differences in tissue density when using microCT to visualize the structure of soft tissue. Our contrast agent, osmium tetroxide^22^, is commonly used because it stains lung tissue intensely and homogeneously penetrates the tissue. Osmium tetroxide preferentially binds to lipids^23^, but when combined with uranyl acetate it effectively illuminates collagen scaffolds in microCT scans^24^. Previous studies that have employed an ex vivo techniques with microCT to image lung tissue have represented tissue density on an 8-bit grayscale, combining it with volume to provide a composite metric of fibrosis^25^. We converted our 16-bit grayscale intensity values of x-ray attenuation to the Hounsfield Unit (HU) scale because it is calibrated to the density of water, making it possible to quantitatively compare our samples to each other. Even without the use of contrast, however, the HU values obtained with a cone-beam microCT system are not equivalent to those obtained with clinical CT machines, even though linearly correlate with them^26,27^. Therefore, it stands to reason that the exaggerated increase in tissue density that we found also exaggerated our calculated increases in cell density.

Overall, we discovered very minimal differences in structural metrics of the control and bleomycin-induced fibrosis microCT scans such as tissue volume (Table 2) and airspace size (Fig. 2). Distributions of airspace volumes were fit best with sum of two exponentials (Table 2). Both a power law and a single exponential fit reasonably well but had unevenly distributed residuals. In 2D, distributions of airspace areas been shown to follow a power law in lungs fixed at 30 cmH_2_O^28^. The two exponents we found in the present study were an order of magnitude apart, implying the existence of two distinct structures having different spatial scales. MicroCT is not able to reliably resolve alveolar septal edges, so our airspace distributions may not have reflected individual alveoli. We therefore surmise that alveolar ducts and bronchioles are the two different structures responsible for the bi-exponential decay in airspace size we found. The bleomycin-induced fibrosis lungs had larger airspaces compared to controls, which has been reported previously^29^. This presumably occurs because as airspaces collapse, they exert traction forces that pull on the remaining open airspaces, causing them to dilate. Nevertheless, we may not have seen the most dramatic differences in parenchymal structure because of the short duration of fibrosis development (21 days) and because the lungs were fixed at a relatively low inflation that, while physiologically relevant, may not have accentuated the volumetric differences between conditions (Fig. 3).

Our previous ABM study simulated two types of agents representing generalized profibrotic and antifibrotic cells that randomly explored and modified a 2D matrix^12^. In expanding this model to 3D, we opted to simulate the behavior of the antifibrotic agents only indirectly for practical reasons since simulating a random walk in 3D is computationally expensive. To compensate, we included global decay term that was applied to all unvisited voxels after 10 time steps. A Bernoulli variable, i.e., a coin flip, was added to make the decay stochastic and thus less aggressive. Previous computational models have used idealize parenchymal microarchitectures for simulation lung disease progression and micromechanics. These architectures have included ensembles of either hexagons, cubes, or truncated octahedra^30–36^. Such models are useful to a point, but our work shows they would have benefitted from inclusion of more realistic architectures. The insights we have obtained thus add to a growing body of literature in the field ^37,38^.

Our study has a number of limitations largely related to the simplifications we made in constructing the ABM. For example, we did not include the effects of substrate biomechanical properties on cell movement. This was done to allow us to focus primarily on the effects of tissue architecture, but the inclusion of more detail about the control of cell movement might have provided more insight into the pathogenesis of IPF. Future developments of the model might also include the addition of mechanical differences between healthy and fibrotic lungs at higher pressures, and the effects this would have had on agent behavior. Such additions might provide insight into the elusive mechanisms of IPF pathogenesis as well as characteristic morphologic features such as peripheral honeycombing^1,2,39^.

In conclusion, we have developed an agent-based model based on microCT lung architecture that maintains a realistic parenchymal structure and density. Increasing agent density in the model was sufficient to produce the characteristic distribution of tissue density seen in fibrotic lungs. We also found that the architecture of actual lung parenchyma, as opposed to an artificial regular lattice structure, widens the distribution of tissue density by forcing agent movement to be more diffuse.

## Methods

### Mouse model of fibrosis

Eight-week-old female C57/BL6 mice (Jackson Laboratories) were studied under the University of Colorado Denver Institutional Animal Care and Use Committee (IACUC) approved protocol #00230. Fibrosis was induced by intratracheal instillation of 4U/kg bleomycin (Meitheal Pharma, Chicago, IL) 21 days prior to experimentation in the BLM group. Controls (CTL) were aged matched. On the day of ventilation, the animals were anesthetized with an intraperitoneal (IP) injection of 100mg/kg ketamine, 8mg/kg xylazine, and 2.5mg/kg acepromazine. Once a deep plane of anesthesia was reached, the mice were tracheostomized with a blunted thin-wall 18-gauge metal cannula and affixed to the FlexiVent rodent ventilator (SCIREQ, Montreal, QC, Canada). Spontaneous breathing efforts were suppressed *via* 0.8 mg/kg pancuronium bromide administered at the onset of ventilation. If necessary, alternating doses of 50 mg/kg ketamine and 50 mg/kg ketamine with 8 mg/kg xylazine were administered IP at 30 min intervals.

### Assessment of lung function

The mice were stabilized for 6 minutes of ventilation at a tidal volume (Vt) = 10 mL/kg, respiratory rate (RR) = 150 breaths/min, positive end expiratory pressure (PEEP) = 3 cmH _2_O, and inspiratory:expiratory ratio (I:E) = 1:1.5. Recruitment maneuvers (RMs) were performed at two-minute intervals. Ventilation was changed to Vt = 6 mL/kg with RR=250 and I:E = 1:1.5 and then Lung function was then assessed. First, RM was performed and then a quasi-static pressure volume loop (PV) was recorded to determine the delivered volume from 0 to 30 cmH _2_O (inspiratory capacity, IC), the slope at 5 cmH_2_O on the expiratory limb (quasi-static compliance, Cst), and hysteresis area (A). We then set PEEP = 9 cmH_2_O, performed another RM, and recorded 6 forced oscillation (FOT) measurements at 20 sec increments which were fit to the constant phase model to determine elastance (H), tissue damping (G), and Newtonian resistance (Rn). The RM and FOT measurements were repeated at PEEP = 6, 3, and 0 cmH_2_O. Finally, a 2^nd^ quasi-static PV loop was recorded.

### Perfusion fixation

After lung mechanics measurements, mice were kept on baseline ventilation (10mL kg, 150 breaths/min, PEEP=3 cmH2O). A bilateral thoracotomy was performed, and the pulmonary vasculature was flushed with a solution of 5,000U/mL heparin and 0.5% sodium nitrate in 0.9% Saline solution for about 5 minutes at a pressure of 40cmH2O until the outflow was clear (typically 5 mL of perfusate). Three consecutive deep inflations were applied that ramped lung pressure up to 30 cmH2O, held for 3 seconds, and returned to PEEP=5 cmH2O. Pressure was then increased to 30 cmH2O and decreased to 5 cmH2O and held while the trachea was ligated. Approximately 5mL of the fixative solution (4% paraformaldehyde and 2% glutaraldehyde in 0.15M HEPES buffer) was then perfused through the right ventricle at 40cmH2O. The lungs were then excised, and immersion fixed for at least 24 hours.

### Lung tissue preparation

The heart and remaining connective tissue were removed from the fixed lungs and the lung volume was measured via volume displacement of distilled water. The lungs were embedded in 3% agar, sliced in the sagittal plane into 2mm slabs, returned to 0.15M HEPES buffer, and vacuum degassed for 3 hours. The slices were then rinsed 3×5 minutes with 0.1M cacodylate buffer and degassed for 10 minutes during the last rinse. The cacodylate buffer was pipetted off and replaced with 1% osmium tetroxide, ensuring the lung slices were completely submerged. The plate was sealed and incubated on ice for 2 hours. Then the osmium tetroxide was removed and rinsed 4×5 minutes with cacodylate buffer. The cacodylate buffer was replaced with distilled water and degassed for 40 minutes prior to overnight immersion in 2% uranyl acetate at 4°C. The slabs were then rinsed 4×5 minutes with distilled water, once with HEPES buffer, and degassed again. The slices were dehydrated in a graded 70% to 100% acetone series for 1.5 hours total and then placed in a 50/50 solution of 100% acetone and Technovit 7100 base liquid solution for 2 hours of infiltration. This solution was replaced by the Technovit 7100 base liquid solution and infiltrated overnight at 4°C. Finally, the lungs were embedded in Technovit 7100 and the glycol methacrylate (GMA) lung blocks were stored in a desiccator until analysis.

### MicroCT and image processing

MicroCT scans were performed with a Bruker 1173 SkyScan. GMA blocks of lung tissue were cut up into sections with a dimension roughly 1cm by 1/2cm by 1/2cm. Samples were adhered to the stage with tack such that the longer dimension was vertical. The sample was moved as close as possible to the X-ray source to allow a resolution of 4.9-micron voxels (volumetric pixels). The X-ray tube voltage was set to 30kV, and the current was set to 160μA. Samples were rotated 360 degrees in 1/10° increments, receiving a 3.5-3.65 second exposure each time. The resultant image was an average of 3 frames.

Reconstruction of the three-dimensional sample was performed with NRecon. The first local minimum in the distribution of grayscale intensity was used as the threshold to exclude the GMA block material. The reconstructed images were exported in the DICOM file format. Further processing was performed in MATLAB. The DICOM files were imported and converted to a 3D matrix. A top hat filter was applied using a sphere with a diameter of 7 voxels as the reference object. Then the largest connected component was selected as the lung structure, considering connectedness of a voxel by another voxel touching its face, edge or corner, excluding all other voxels. For each sample, a region of interest (ROI) was selected such that the 108 x 108 x 216 voxel cuboid did not intersect bronchi or the pleural surface and mostly contained alveoli with a few bronchioles. For the ABM, these cuboids were divided up into 27 x 27 x 27 voxel cubes to be run independently. Disconnected clusters of voxels containing five or fewer voxels were removed. Grayscale intensity values were linearly transformed to the Hounsfield units so that they were calibrated to the same relative scale; we refer to these as contrast-enhanced Hounsfield units (ceHU). The ceHUs were scaled down such that the mean value of the control structures’ ceHUs equaled β.

### Lattice architecture

As a control structure, cubic lattices of the same size as the lung matrices were generated. The lattices were required to alternate between planes of filled and empty spaces to represent tissue and airspaces in all three directions. The number of layers of filled planes and empty planes was kept constant. The lattices are required to be symmetric in each x-, y-, and z-direction for tessellation. Lastly, lattices required a minimum volume fraction of 0.2. There were 45 unique layouts that fit these criteria, however, for comparisons to lung tissue we only used lattices whose volume fraction was between the minimum and maximum of the lung matrices, 0.44 and 0.84 respectively, resulting in 15 unique layouts used as structural controls (outlined points in Sup. Fig. 1). All tissue values were initialized to our β parameter.

### Structural analysis

Distribution of airspaces from lung micro-CT scans of the control and bleomycin-induced fibrosis mice were used to compare structures. Sizes of airspace areas were counted plane-by-plane from the post-processed matrix of lung tissue. This process was repeated for each of the xy-, xz, and yz-planes and pooled together as a distribution of airspace areas. This process was done for each ROI and whole lung tissue that the ROI is a subsection of. Volume fraction was defined as the ratio of number of filled voxels to total number of voxels.

### Random walks

Movement for our ABM was simulated as a discrete Markov chain with the MATLAB function, `dtmc`. The Markov chain was generated with the adjacency matrix of the structure matrix where locations were not considered connected to themselves. Agents are only allowed to move 1 space at a time on filled voxels representing discretized lung tissue. Our model used a random walk that allowed agents to move to any of the up to 26 adjacent, filled voxels with equal probabilities.

### Agent-based model (ABM)

Our ABM was an extension on a previous ABM of tissue maintenance^12^, described by the following equation:

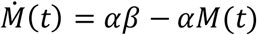

For the current model, agents again added a fixed amount of tissue with each step, which is tallied every 8 steps. Every unvisited location within the 10-step time frame would be decreased by a fraction of the tissue at those locations. These actions are described by the following equation:

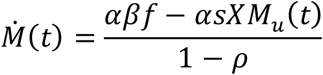

where *f* is the frequency that a location has been visited, *s* is the number of steps taken, *X*∼Bern(1/2), *M_u_* is the tissue value of the unvisited locations, and *ρ* is our agent density, which is the ratio of number of agents to number of voxels containing tissue. These steps were repeated for 10,000-time steps. This model was run on an ROI from all three animals as well as lattices that correspond to the same overall matrix size. These were done with a random walk on each architecture at the following agent densities: 0.1365, 0.175, and 0.20.

### Statistical analysis

All statistical tests were performed with MATLAB 2022a. Parameters from FOT (Table 6) were tested via a two-sample *t*-test for unequal variances^40^. A p-value < 0.05 was considered significant. Nonlinear least squares regression of the sum of two exponentials (Table 2) was fitted to the following equation:

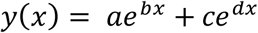

**Table 6:** p-values for Welch’s t-test between control and bleomycin-induced fibrosis of lung mechanics measurements at various PEEP.

|  | PEEP=0 | PEEP=3 | PEEP=6 | PEEP=9 |
| --- | --- | --- | --- | --- |
| H | p=0.003 | p=0.002 | p=0.004 | p=0.011 |
| G | p=0.004 | p=0.011 | p=0.007 | p=0.008 |
| Rn | p=0.035 | p=0.015 | p=0.055 | p=0.171 |
| G/H | p=0.345 | p=0.647 | p=0.906 | p=0.859 |

The power law fit (Table 4) is described by the following equation:

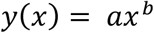

## Acknowledgements

This study was supported by NIH grants T32 HL-076122, U01 HL139466, R01HL151630, and NSF 2225554. Computations were performed, in part, on the Vermont Advanced Computing Center. We would like to thank J. Matthew Mahoney for advice on the ABM and Lars Knudsen for guidance on microCT.

## Author contributions statement

DTC coded the model and ran the analysis, performed the CT, and drafted the manuscript. AS performed mouse experiment, lung function testing, and processed lung tissue. DTC, VM, JKH, BJS, BS and JHTB developed the modeling methodology, interpreted the results, and edited the manuscript. All authors approved the final submission.

---

**Sup. Figure 1:**
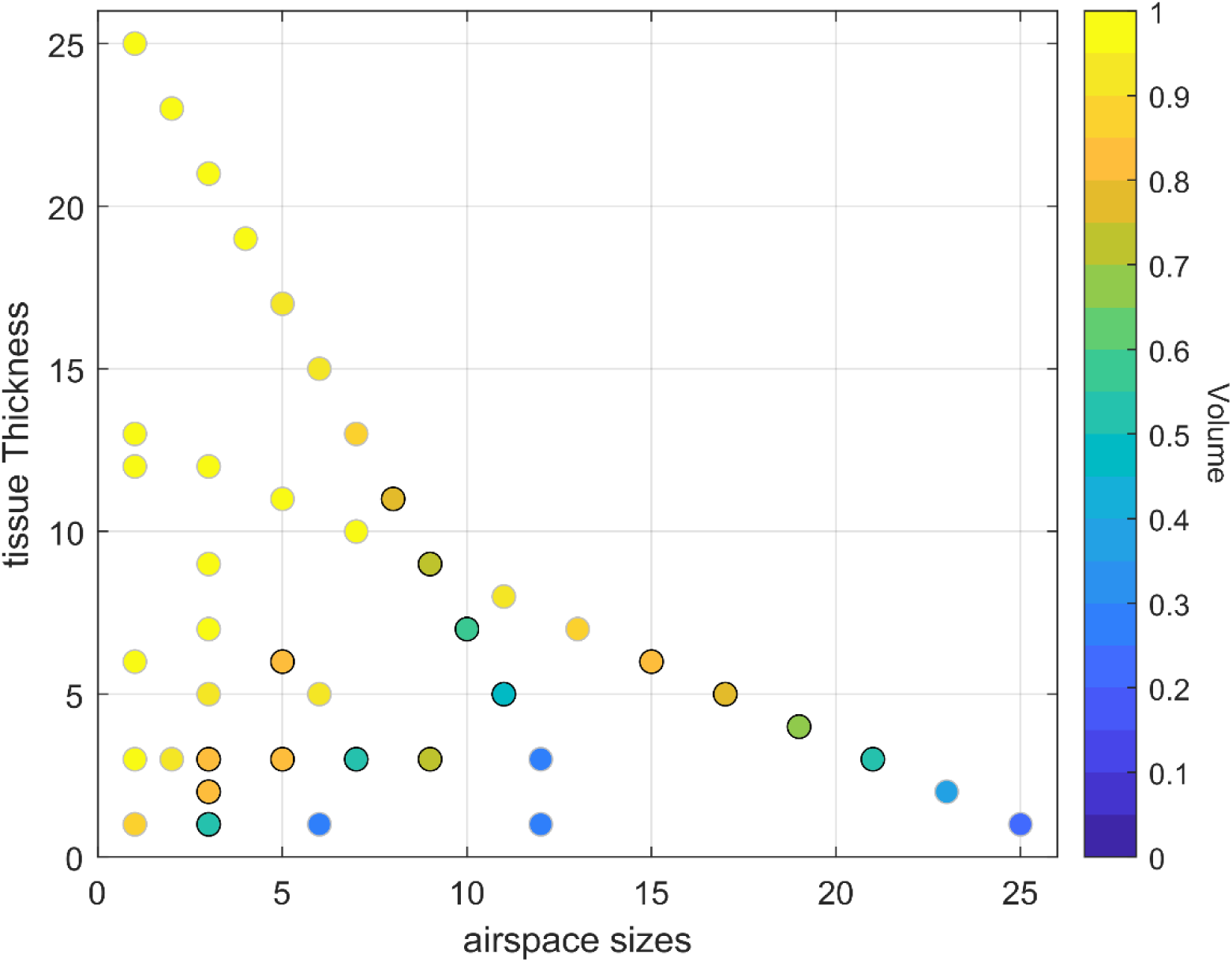
Plot of lattice architecture patterns. Tissue and airspace sizes were alternated up to size 27. Color represents the volume of tissue those patterns have. Dark outlined points are architectures used to run the ABM i.e. has volume between 0.44 and 0.84.

**Sup. Figure 2:**
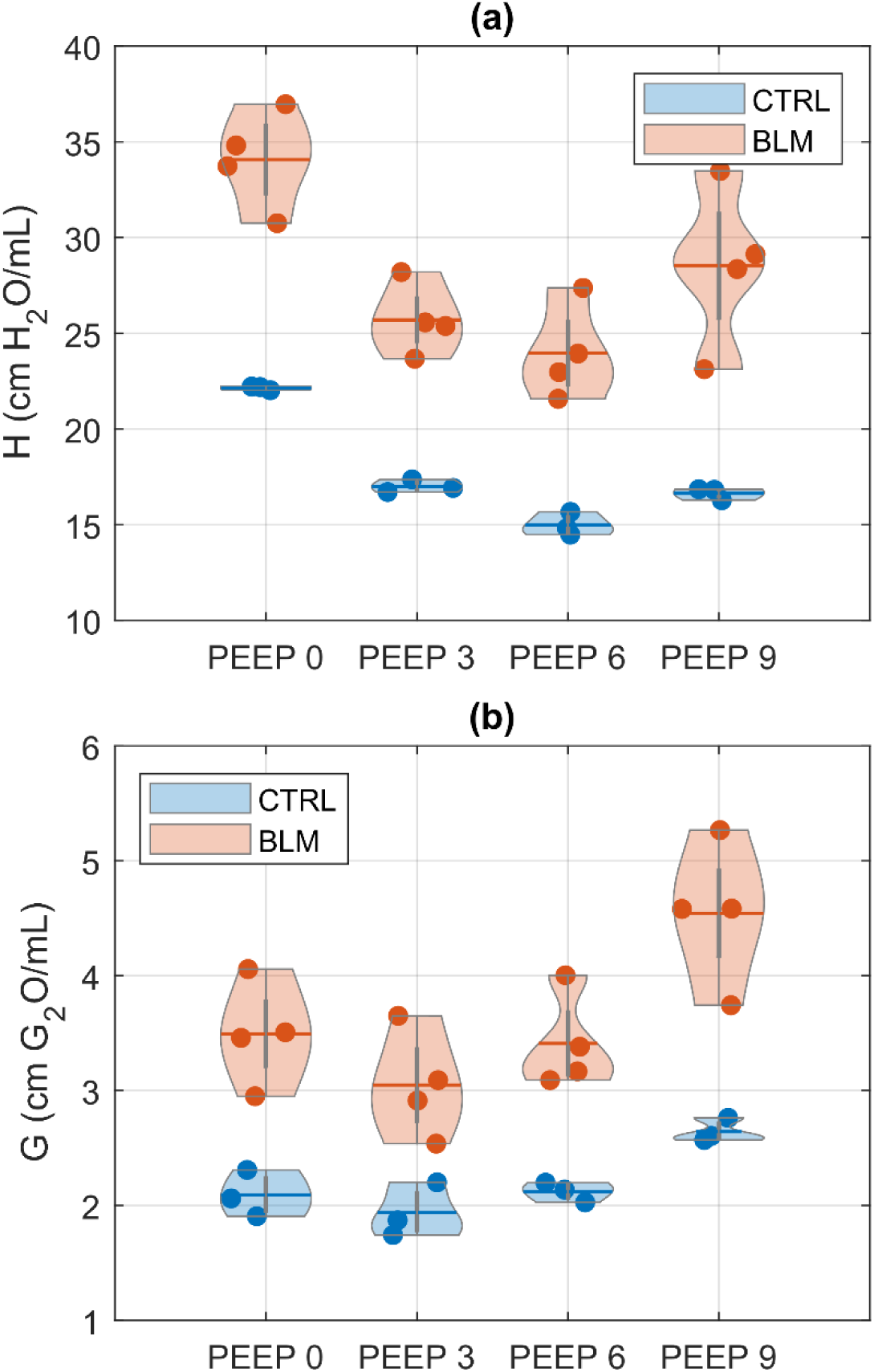
Parameters from forced oscillation technique for control (n=3) and bleomycin-induced fibrosis mice (n=4). As a function of PEEP, **(a)** H is the elastance of lung tissue and **(b)** G is the tissue damping. Results from Welch’s t-test for G and H are in Table 6.

**Sup. Figure 3:**
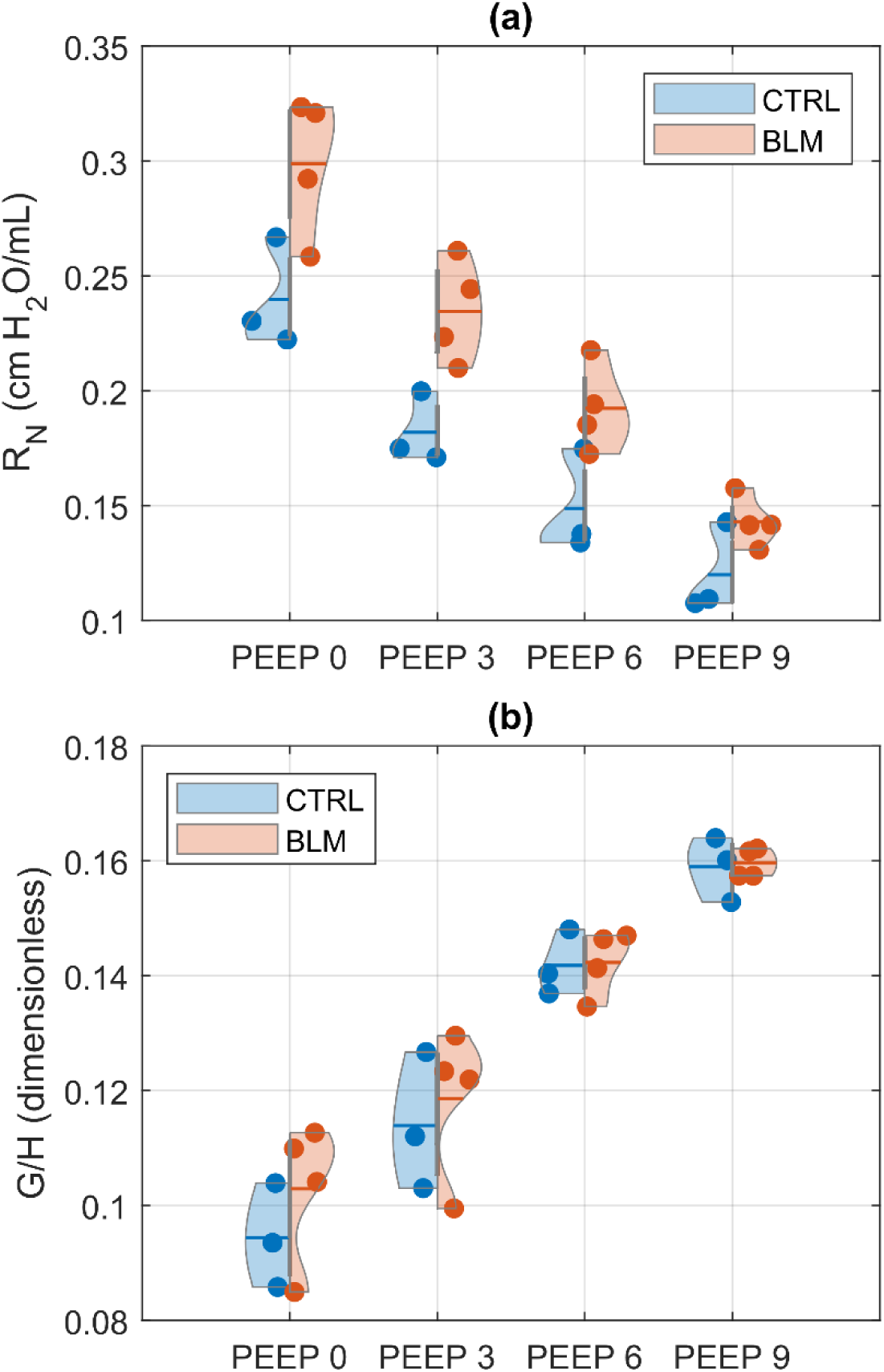
Parameters from forced oscillation technique for control (n=3) and bleomycin-induced fibrosis mice (n=4). As a function of PEEP, **(a)** Rn is the airway impedance and **(b)** G/H is the hysteresivity. Results from Welch’s t-test for Rn and G/H are in Table 6.

**Sup. Figure 4:**
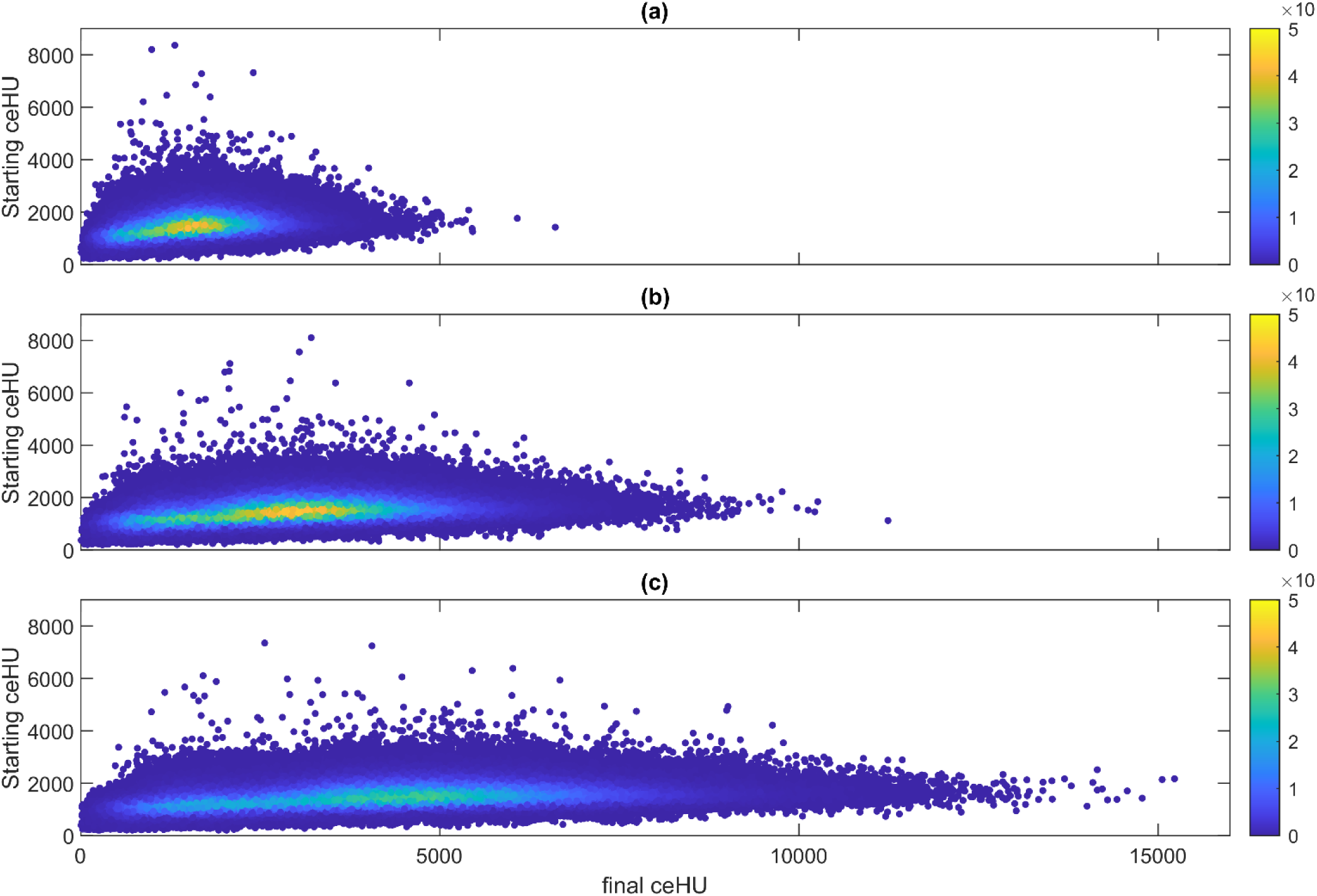
Comparison of ceHU at time step 0 and 10,000 for lung architecture. (a) represents the results from agent density of 13.65%, (b) is from agent density of 17.5%, and (c) is from agent density of 20%. Color bar represents density.

